# A simple method for mapping the location of cross-β forming regions within protein domains of low sequence complexity

**DOI:** 10.1101/2025.02.13.638120

**Authors:** Jinge Gu, Xiaoming Zhou, Lillian Sutherland, Glen Liszczak, Steven L. McKnight

## Abstract

Protein domains of low sequence complexity are unable to fold into stable, three-dimensional structures. In test tube studies, these unusual polypeptide regions can self-associate in a manner causing phase separation from aqueous solution. This form of protein:protein interaction has been implicated in numerous examples of dynamic morphological organization within eukaryotic cells. In several cases, the basis for low complexity domain (LCD) self-association and phase separation has been traced to the formation of labile cross-β structures. The primary energetic force favoring formation of these transient and reversible structures is enabled by polypeptide backbone interactions. Short, contiguous networks of peptide backbone amino groups and carbonyl oxygens are zippered together intermolecularly by hydrogen bonding as described by Linus Pauling seven decades ago. Here we describe a simple, molecular biological method useful for the identification of localized, self-associating regions within larger protein domains of low sequence complexity.

**Significance:** This study describes a molecular biological method for analyzing protein domains of low sequence complexity in search of segments that mediate self-association and consequent phase separation both *in vitro* and *in vivo*. Small regions allowing for self-association correspond to sequences that specify the formation of labile cross-β structural order. When juxtaposed to the C-terminus of GFP, cross-β prone regions suppress fluorescence. A tiled scan of overlapping fragments of the low complexity domain (LCD) of the TDP-43 RNA binding protein pinpointed an evolutionarily conserved sequence of twenty amino acids essential for self-association, phase separation and the formation of nuclear speckles. The screening method described herein should be useful for the analysis of any LCD believed to function via homotypic self-association.

## Introduction

Most proteins encoded by the genes of all life forms function by adopting a unique conformation of structural order. Three-dimensional structural order is encoded by the linear sequence of amino acids of a protein. This biological truth was confirmed by the unfolding and folding experiments of Anfinsen and colleagues using ribonuclease (1).

Unusual protein sequences incapable of folding into stable structures began to be described three to four decades ago. Early examples of these strange proteins were reported for the activation domains of the Gal4 and VP16 transcription factor (2–4). The amino acid sequences of the Gal4 and VP16 transcriptional activation domains were unusual in being composed of a limited number of the twenty residues utilized for the folding of normal proteins. Despite being able to perform a distinct biological function evolutionarily crafted to stimulate transcription, these unusual protein domains were incapable of folding into stable molecular structures.

In the ensuing decades it became clear that the proteomes of eukaryotic organisms contain thousands of these protein domains of low sequence complexity. Such regions have variously been referred to as low complexity domains (LCDs), intrinsically disordered domains (IDRs), or prion-like domains (PLDs). Extensive studies have confirmed that most of these low complexity domains are either incapable of folding into stable, three-dimensional structure, or do so conditionally only upon interaction with other, structurally ordered macromolecules (5, 6). Upwards of a quarter of all eukaryotic proteins, or extensive sub-segments thereof, are now recognized to be of low sequence complexity.

How do these unusual proteins perform their biological function? Independent studies focused on the phenylalanine-rich LCD of the NUP98 nucleoporin protein, and the tyrosine-rich LCD of the fused-in-sarcoma (FUS) RNA-binding protein, reported the formation of phase-separated hydrogels (7–11). Both studies described a form of protein self-association dependent upon aromatic amino acids. Both studies likewise gave evidence that LCD self-association involves the formation of polypeptide backbone-mediated cross-β interactions. These unexpected observations of LCD phase separation raised the possibility that the nucleoporin and FUS proteins might execute their biological function by forming some form of weakly-tethered, homotypic interactions.

Subsequent studies have functionally mapped the cross-β segments responsible for mediating a number of distinct examples of LCD self-association (12–18). The small regions specifying self-association have, in several cases, been found to co-localize with mutations causative of human disease. Disease-causing mutations proximal to cross-β forming regions within LCDs specify the synthesis of altered proteins that aberrantly strengthen the structurally ordered state (13, 15, 16). Such observations give evidence that the biological utility of this form of protein function requires a type of weak intermolecular self-association poised at the balance of thermodynamic equilibrium. Various forms of post-translational modification also map within the cross-β forming regions of LCDs and serve to regulate the balance of self-association (12, 14, 15, 24–26).

Perhaps the clearest description of how LCD self-association abets the dynamics of cell morphology has come from studies of the intrinsically disordered head domains of intermediate filament (IF) proteins. IF head domains self-associate in the form of weakly annealed cross-β structures (15, 16, 27, 28). A beautiful cryo-EM tomography study of assembled vimentin IFs has revealed the periodically phased coalescence of 40 head domains into a luminal cavity (29). It is within this cavity that IF head domains self-adhere to facilitate filament assembly. The orthogonal experimental findings leading to this understanding of IF architecture offer a refreshingly simple description of how head domain phosphorylation triggers filament disassembly and the rounding up of cells during mitosis (30).

A comprehensive understanding of how our thousands of LCDs participate in the assembly and regulation of malleable cellular structures will require machine learning-assisted identification of the localized regions within LCDs that mediate transient annealing. Instead of being limited to a handful of examples of tediously-mapped LCD sequences, experimentalists need to provide computational scientists with hundreds of accurately defined cross-β forming regions within LCDs.

To this end we have turned to methods described by Moffet and colleagues who have shown that the amyloidogenic region of the pancreatic islet amyloid polypeptide (IAPP) suppresses fluorescence upon fusion to the C-terminus of green fluorescent protein (31). This method was used to identify mutations within the IAPP that block the formation of amyloid polymers. Expanding from the teachings of Moffet, we describe a technically simple method for mapping the location of the cross-β forming region that mediates self-association, phase separation and biological function of the LCD associated with the TAR DNA-binding protein 43 (TDP-43). Aside from assisting in the development of computational methods required to evolve studies of LCDs from lab bench to computer, the methods described herein will also be of use for the functional analysis of intrinsically disordered protein domains.

## Results

The LCD of the TDP-43 RNA binding protein extends from proline residue 262 to the carboxyl terminus of the protein, ending with methionine residue 414. An evolutionarily-conserved region of roughly 20 amino acids located between residues 321 and 340 has been found to mediate interstrand self-association via the formation of a labile cross-β molecular structure (24). This form of protein self-association is enabled by the zippering of a contiguous network of interstrand hydrogen bonds between peptide backbone amino groups and carbonyl oxygen atoms (16).

The experimental path leading to these conclusions has been slow and tedious. In efforts to develop more facile methods for mapping the small, functionally critical regions of LCDs required for self-association and biological function, we turned to studies published by David Moffet and colleagues who have shown that the pancreatic islet amyloid polypeptide (IAPP) protein suppresses fluorescence upon fusion to the C-terminus of GFP (31). Following the teachings of Moffet, we interrogated the TDP-43 LCD by systematically fusing individual, 30 residue fragments to the C-terminus of GFP (Figure 1A). Fragments overlapping by twenty residues were systematically scanned across the LCD. Plasmids encoding twelve GFP:TDP-43 fusion proteins were introduced into bacterial cells and subjected to conditions allowing for induced protein expression (Materials and Methods).

**Figure 1.**
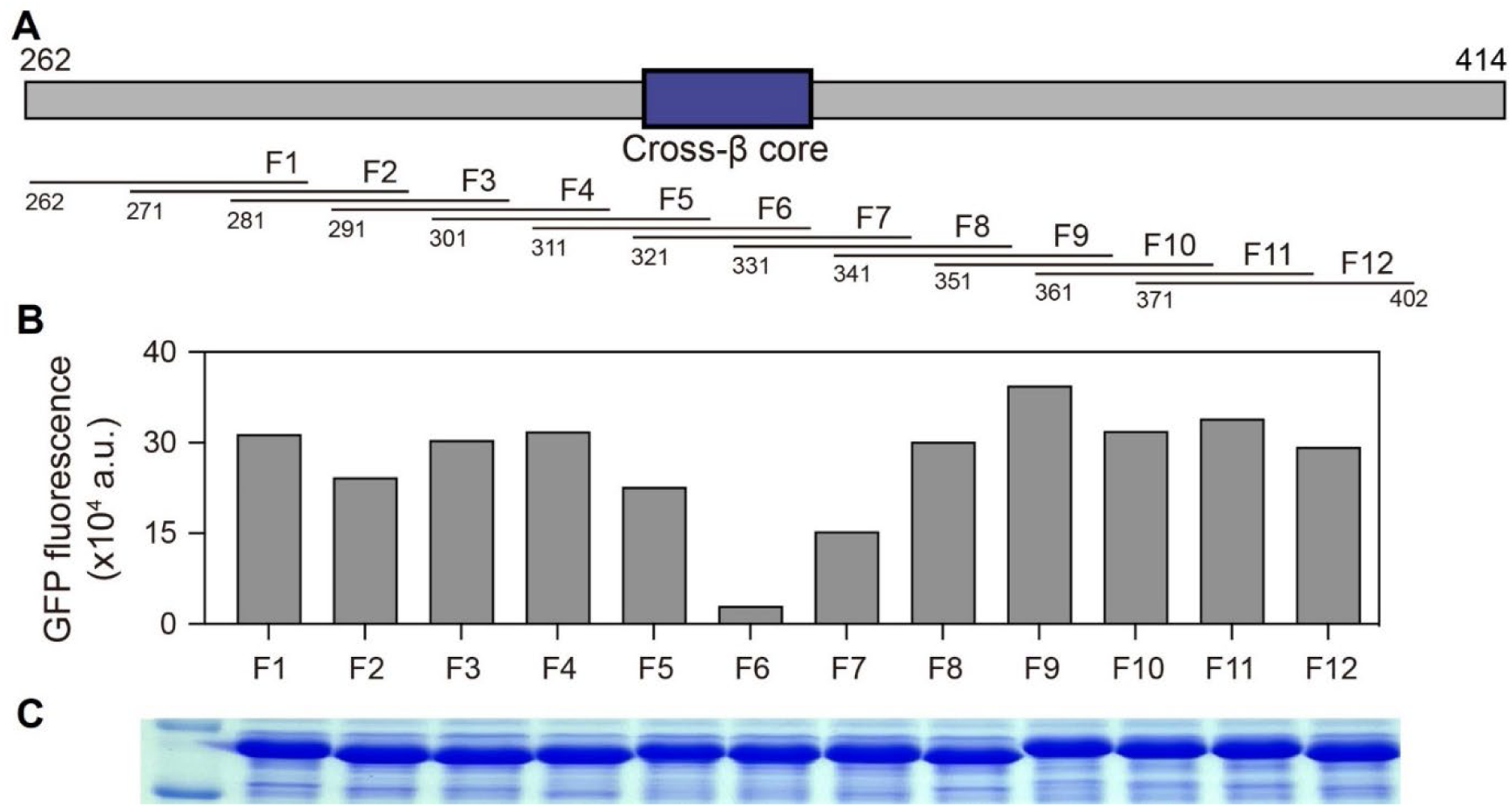
Fusion of the cross-β core of the TDP-43 low complexity domain onto the carboxyl terminus of GFP impedes fluorescence. (A) Schematic diagram of the low complexity domain of TDP-43 spanning residues 262 through 414. Location of the cross-β core responsible for self-association and phase separation is shown as a blue box. Overlapping segments of 30 residues were individually appended onto the carboxyl terminus of GFP and expressed in bacterial cells (Materials and Methods). (B) Spectrophotometric measurements of GFP fluorescence of soluble lysates prepared from bacteria induced to express GFP fusion proteins. (C) Coomassie-stained SDS polyacrylamide gel used to size and visualize bacterially-expressed GFP fusion proteins.

Cell lysates were evaluated by SDS polyacrylamide gel electrophoresis (SDS-PAGE) to monitor protein expression, and spectrophotometry to measure GFP fluorescence (Figure 1B and 1C). As shown in Figure 1C, all twelve of the GFP fusion proteins were expressed at equivalent levels as determined by SDS-PAGE analysis of bacterial lysates. Roughly equivalent levels of GFP fluorescence were observed for ten of the fusion proteins (Figure 1B). By contrast, the protein linking residues 311-342 (fragment 6, F6) of the TDP-43 LCD to the C-terminus of GFP suffered a substantial reduction in fluorescence, and the fusion protein linking GFP to residues 321-351 (fragment 7, F7) exhibited a modest reduction in GFP fluorescence.

The twenty-residue overlap of the latter two segments corresponds to residues 321-342 of the TDP-43 LCD. As reported in earlier studies, this span of twenty amino acids is the most evolutionarily conserved region of the TDP-43 LCD (24). Likewise, this GFP-inhibiting region corresponds with the protein sequences essential for formation of both phase-separated, liquid-like droplets and labile cross-β polymers (16).

### A randomized screen to identify GFP-inhibiting fragments within the TDP-43 LCD

Observations shown in Figure 1 give evidence that fusion of either of two, 30-residue segments of the TDP-43 LCD to the carboxyl terminus of GFP interferes with fluorescence. Reasoning that this might form the basis for a functional screen in bacteria, a plasmid library was assembled wherein synthetic oligonucleotides corresponding to 30-residue segments of the TDP-43 LCD were shotgun cloned downstream of the coding sequence of GFP (Figure 2A). Competent BL21(DE3) bacterial cells were transformed with the library and spread on agarose plates supplemented with IPTG. Visual inspection of the culture plates under UV light revealed a mixture of bright green and dim green colonies (Figure 2B). 370 colonies, scored as being either bright or dim green in color, were picked and sequenced.

**Figure 2.**
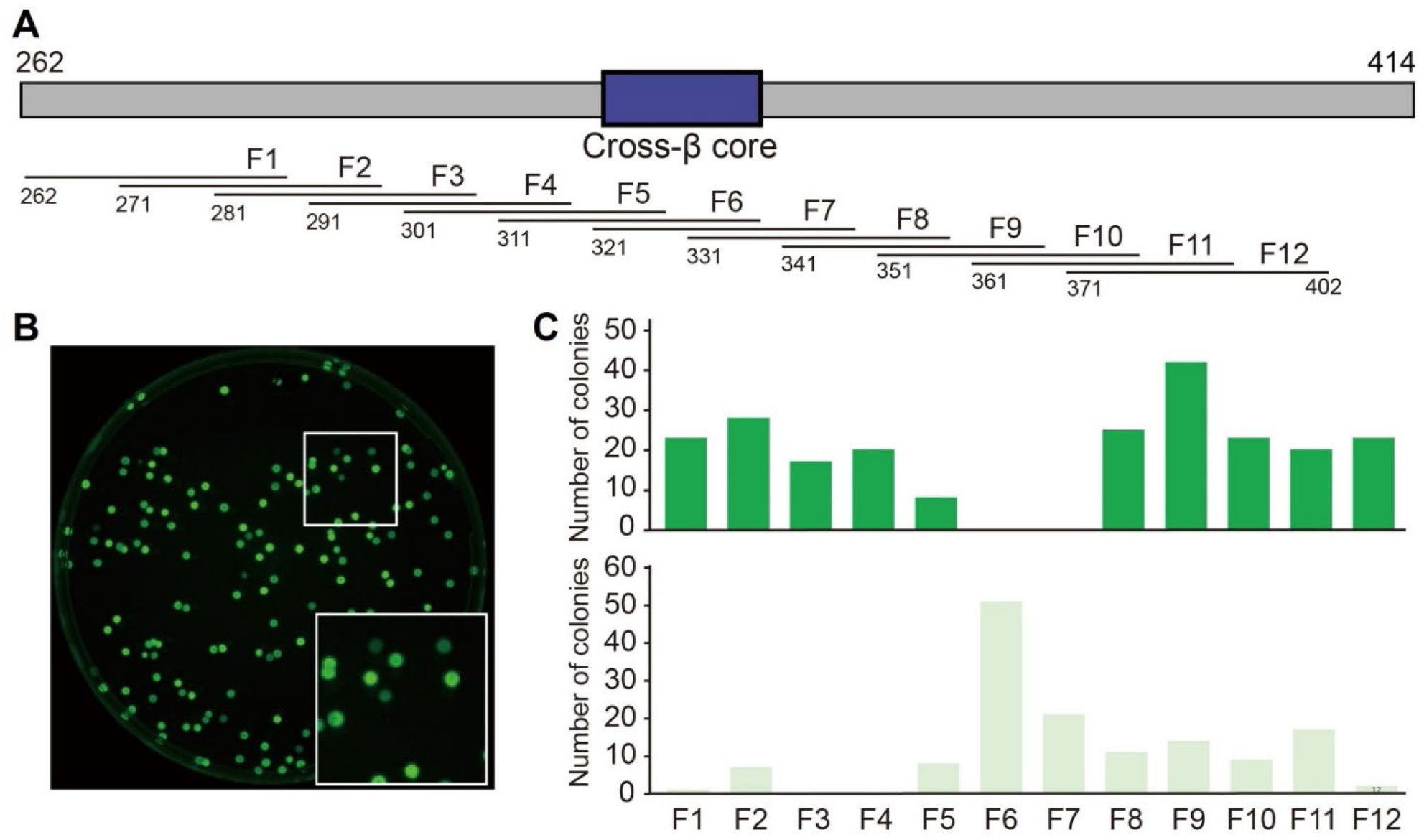
Analysis of bacterial colonies expressing GFP fused to 30 residue fragments of the TDP-43 low complexity domain. (A) Schematic diagram of the low complexity domain of TDP-43 spanning residues 262 through 414. Location of the cross-β forming region responsible for self-association and phase separation is shown as a blue box. (B) Twelve, double-stranded synthetic oligonucleotides encoding 30 residue fragments of the TDP-43 low complexity domain were mixed and shotgun cloned immediately downstream from the C-terminus of GFP. The plasmid library was transformed into bacterial cells that were grown on agarose plates in the presence of 40 μM IPTG at 37 °C to induce GFP expression. Colonies scored as either bright green or dim green were picked and sequenced from plates illuminated by UV light. (C) Quantification of bright green (above) or dim green (below) colonies containing designated fragments of the TDP-43 low complexity domain.

The histograms of Figure 2C show that no bright green colonies were found to contain sequences encoding the evolutionarily conserved, cross-β core of the TDP-43 LCD (as defined by oligonucleotides encoding residues 312-342 or 321-351). All colonies containing these two sequences, hereafter designated as fragments 6 and 7 of the LCD, had been scored as being dim green. Of the remaining, 30-residue segments of the TDP-43 LCD, all were found by unbiased screening and sequencing to have been derived either exclusively from bright green colonies, or from a mixture of both bright and dim green colonies.

### Mutational analysis of GFP-inhibiting fragments of the TDP-43 LCD

As a test of the possible involvement of β-strand structural conformation in the inhibition of GFP fluorescence, bacterial expression vectors linking either fragment 6 or fragment 7 of the TDP-43 LCD to the C-terminus of GFP were altered by site-directed mutagenesis. Starting with phenylalanine residue 313 for fragment 6, or alanine residue 321 for fragment 7, pairs of residues separated by a single amino acid were changed to proline to systematically mutagenize each fragment. Proline substitutions were chosen because proline is the sole amino acid that does not contribute an NH group to the polypeptide backbone. As such, it disfavors the formation of annealed β-sheets.

The sequences of these double proline variants are shown in Figure 3A. Each variant plasmid was introduced into bacterial cells that were subsequently exposed to IPTG as a means of inducing expression of the encoded GFP fusion protein (Materials and Methods). SDS-PAGE analysis of bacterial lysates confirmed that all mutational variants were expressed at equivalent levels (Supplemental Data, Figure S1).

**Figure 3.**
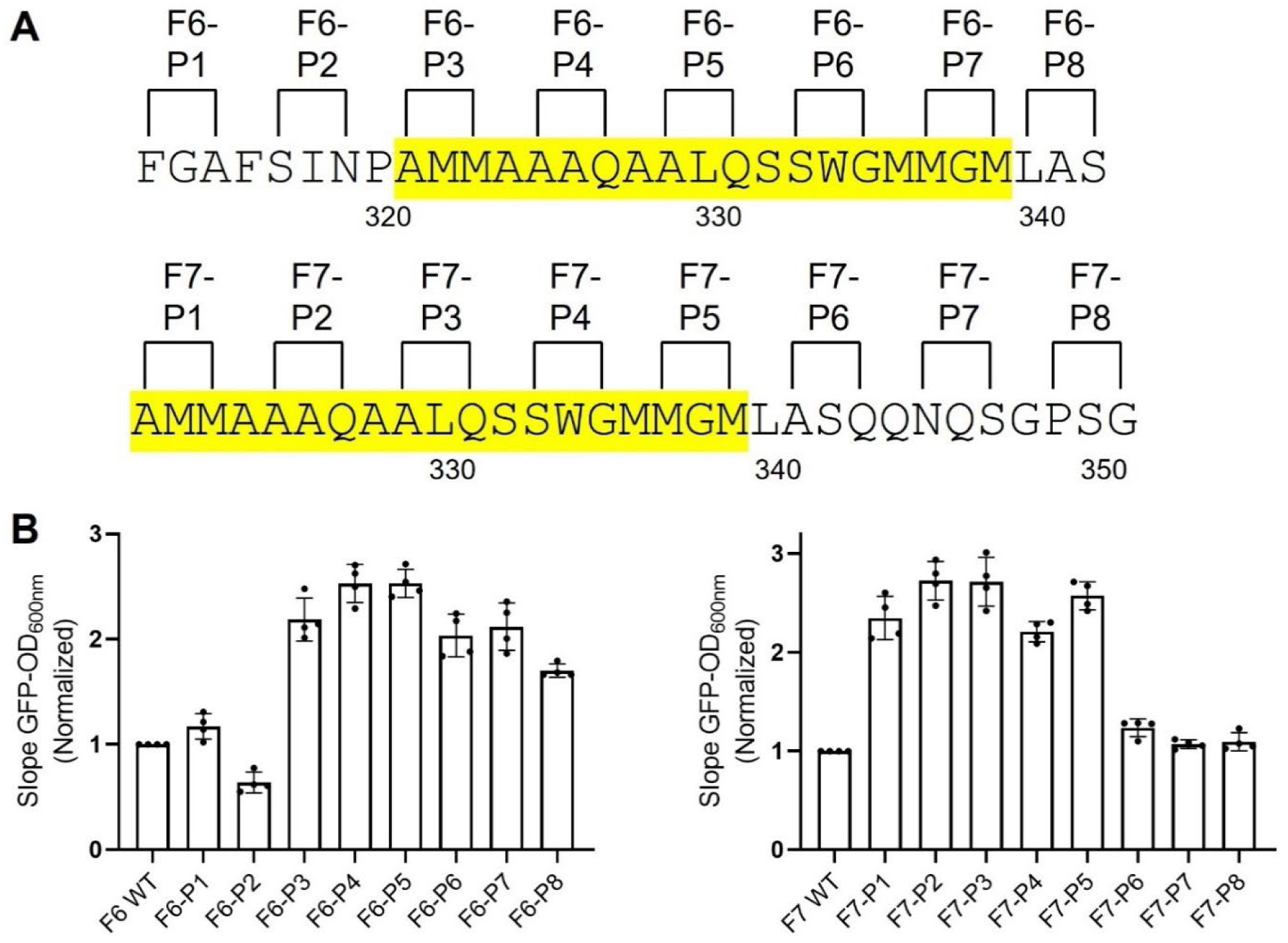
Effects of proline substitutions upon the suppressive activities of fragments 6 and 7 of the TDP-43 low complexity on GFP fluorescence. (A) Sequences of double proline substitution variants of fragments 6 (above) and 7 (below) of the TDP-43 low complexity domain. Sequences highlighted in yellow correspond to cross-β forming region. (B) Measured levels of GFP fluorescence for each double proline variant of fragment 6 (F6, left) and fragment 7 (F7, right) of the TDP-43 low complexity domain as assayed in living bacterial cells. Cultures of bacterial strains expressing each variant were evaluated as a linear function of turbidity (OD600nm) to measure cell density, and intensity of GFP fluorescence (Supplemental Data, Figure S2). Quantitative comparisons of slopes of increasing OD600nm versus increasing GFP intensity, as measured for four independent cultures of each variant, yielded histogram patterns for double proline variants of F6 (left) and F7 (right). Mean ± SD; n = 4.

Four separate samples of IPTG-induced cells were recovered for each variant and measured for GFP fluorescence (Figures 3B and S2). The histograms of Figure 3B (left) reveal the levels of GFP fluorescence observed for each double proline variant introduced into fragment 6 of the TDP-43 LCD. The first two variants, designated F6-P1 and F6-P2, exhibited levels of GFP fluorescence no higher than bacterial cells expressing GFP fused to the native sequence of fragment 6. By contrast, all of the remaining double proline variants of fragment 6 yielded significantly higher levels of GFP fluorescence. We conclude that the latter variants prevent fragment 6 of the TDP-43 LCD from suppressing GFP fluorescence.

The histograms of Figure 3B (right) reveal the distributions of GFP fluorescence observed in bacterial cells expressing each of the double proline variants introduced into fragment 7 of the TDP-43 LCD. The first five variants introduced into fragment 7, upon IPTG-induced expression in bacterial cells, exhibited significant GFP fluorescence. Variants bearing double proline mutations close to the C-terminal side of fragment 7 produced lower levels of GFP fluorescence. The variants designated F7-P6, F7-P7 and F7-P8 exhibited no higher a level of fluorescence than bacterial cells expressing GFP fused to the native sequence of fragment 7. We conclude that variants F7-P1 through F7-P5 interfered with the ability of fragment 7 of the TDP-43 LCD to suppress GFP fluorescence.

The combined data of Figure 3 reveal a contiguous segment of 20 amino acids, spanning alanine residue 321 to leucine residue 340, wherein introduction of double proline variations interfered with the ability of either fragment 6 or fragment 7 of the TDP-43 LCD to suppress GFP fluorescence in bacterial cells. These 20 amino acids represent the most evolutionarily conserved region of the TDP-43 LCD (24). They further correspond to the cross-β forming region essential for formation of phase separated liquid-like droplets and labile cross-β polymers (16).

### Analysis of proline mutations within the TDP-43 LCD upon self-association and phase separation

Having observed double proline variants that either do or do not affect the ability of the TDP-43 LCD to suppress GFP fluorescence in bacterial cells, we next sought to test for possible correlative effects of these mutational variants on two forms of phase separation. Incubation of the isolated LCD of TDP-43 under conditions of neutral pH and physiologically normal monovalent salts leads to the formation of liquid-like droplets (16). Protein samples of either the native LCD of TDP-43, or variants bearing two proline substitutions, were purified and assayed under conditions allowing formation of liquid-like droplets. As shown in Figure 4, six of the double proline variants were compromised in the formation of liquid-like droplets when tested under assay conditions of 150 mM NaCl and pH 6.5. Less severe impediments to droplet formation were observed upon assay at pH 6.8. Under such conditions, droplet formation by the F6-P2, F6-P6 and F7-P6 variants was reduced by only 2-fold relative to the native LCD of TDP-43. The double proline variants observed to impede droplet formation coincided with those that overcame GFP suppression as described in Figure 3. Likewise, double proline variants that did not overcome GFP suppression in live bacterial cells, including F6-P1, F7-P7 and F7-P8, also failed to impede formation of liquid-like droplets.

**Figure 4.**
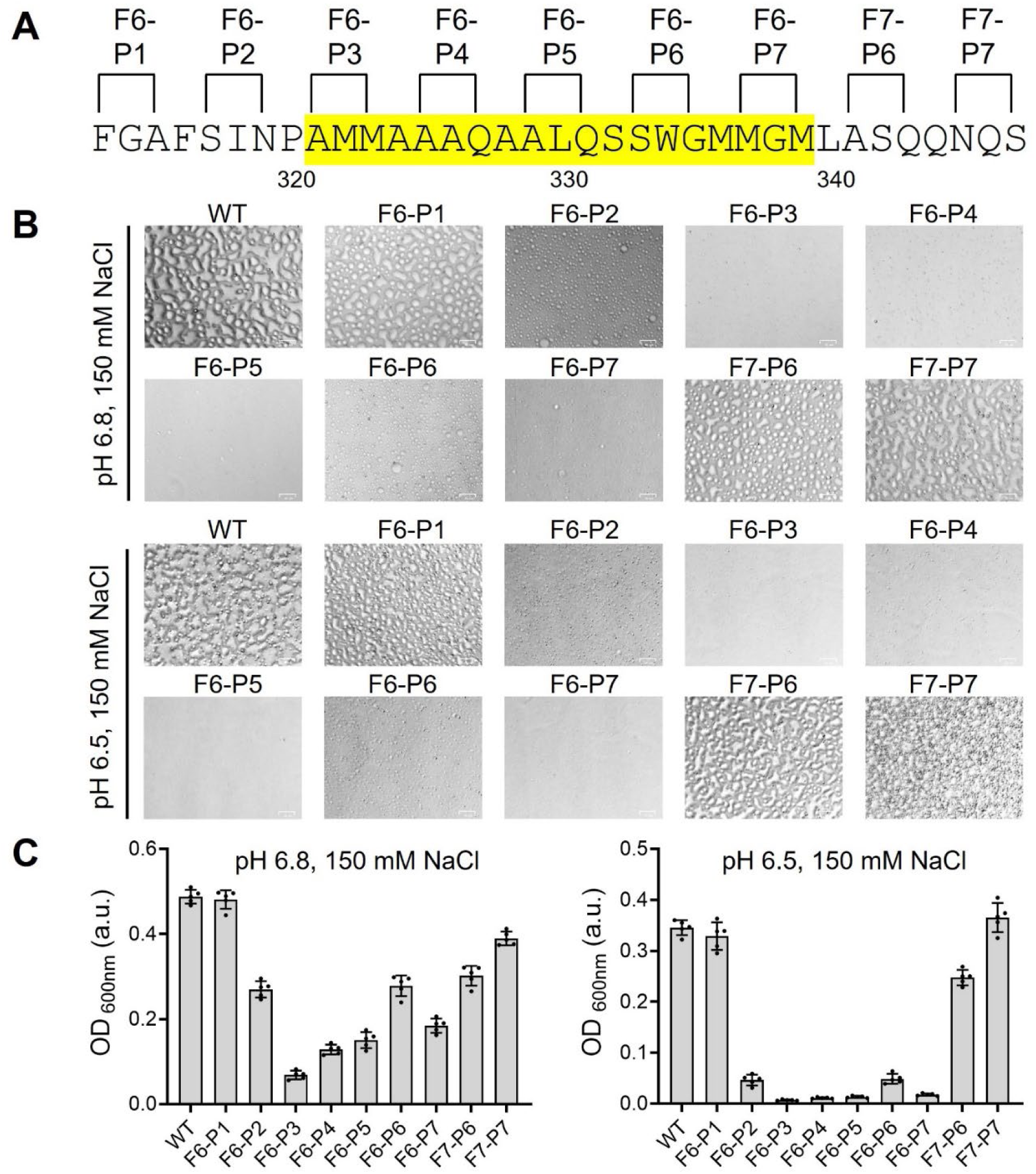
Effects of proline substitutions upon formation of phase separated liquid-like droplets formed by the TDP-43 low complexity domain. (A) Purified samples of the intact low complexity domain of TDP-43, or proline-substituted variants thereof, were tested for phase separation under assay conditions of 150 mM NaCl and pH levels of either 6.8 or 6.5. Locations of proline substitutions are shown on the sequence of the TDP-43 low complexity domain between phenylalanine residue 313 and serine residue 347. Sequence highlighted in yellow corresponds to cross-β forming region. (B) Microscopic images of protein samples incubated under conditions permissive of phase separation revealed the presence, attenuation or absence of liquid-like droplets. Scale bar: 25 μm. (C) Optical density measurements are shown for each protein sample as measured at pH 6.8 (left) or pH 6.5 (right). Mean ± SD; n = 5.

The same protein samples evaluated for formation of phase separated liquid-like droplets were also tested in biochemical assays of the formation of labile cross-β polymers (Figure 5). Samples prepared at a protein concentration of 20 μM were incubated in a buffer of neutral pH and physiological ionic strength in the presence of thioflavin-T (Materials and Methods). Acquisition of enhanced ThT fluorescence was monitored over a period of 30 h. Time-dependent enhancement of ThT fluorescence was readily observed for the native protein, the F6-P1 variant and the F7-P7 variant. Either delayed or intermediate levels of ThT fluorescence were observed for the F6-P2 and F7-P6 variants. Little or no evidence of time-dependent increase in ThT fluorescence was observed for the F6-P3, F6-P4, F6-P5, F6-P6 or F6-P7 variants.

**Figure 5.**
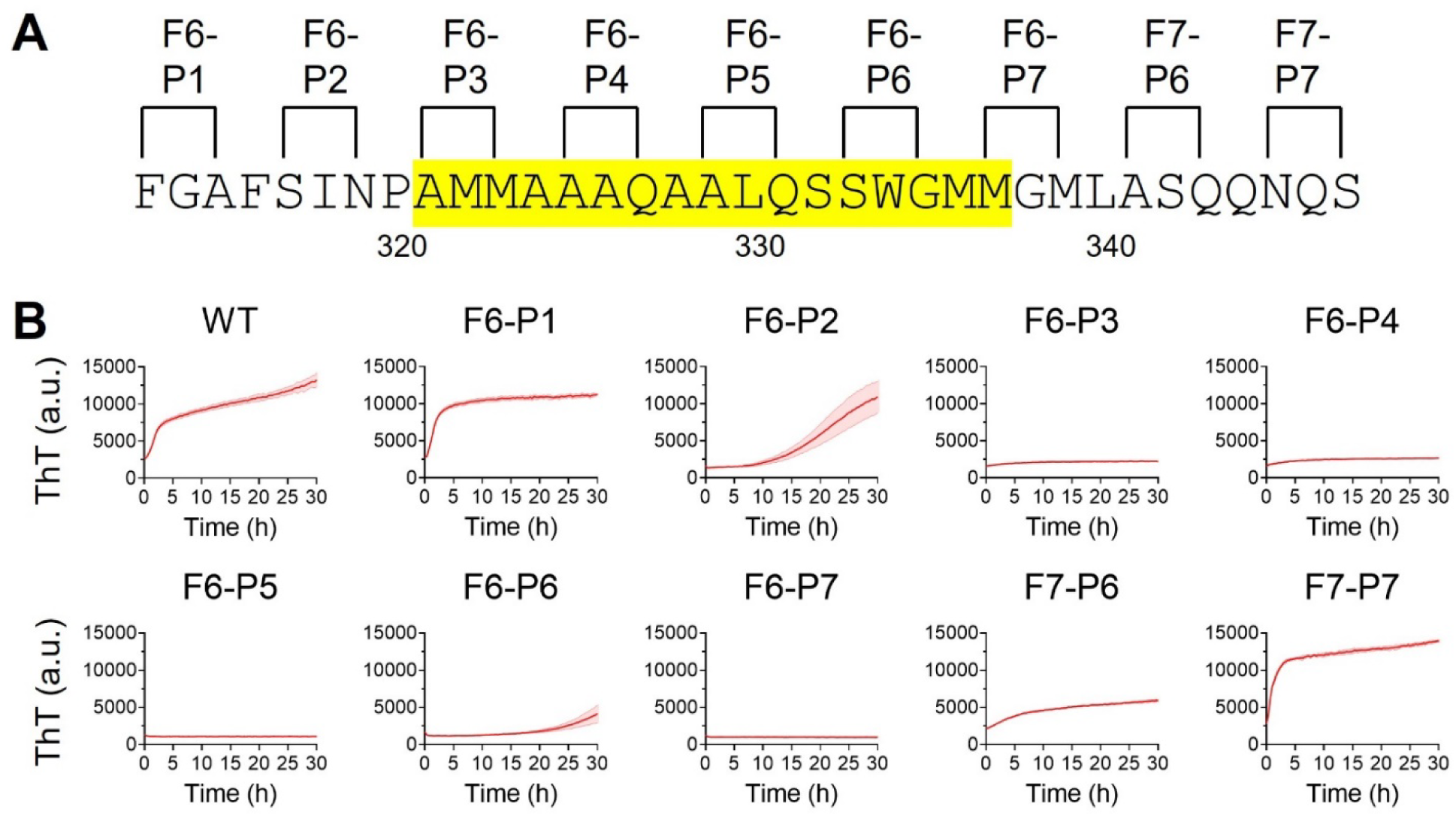
Effects of proline substitutions upon formation of labile cross-β polymers formed by the TDP-43 low complexity domain. (A) Purified samples of the intact low complexity domain of TDP-43, or proline-substituted variants thereof, were tested for the formation of cross-β polymers as revealed by the enhanced fluorescence of thioflavin-T (ThT). Locations of proline substitutions are shown on the sequence of the TDP-43 low complexity domain between phenylalanine residue 313 and serine residue 347. Sequence highlighted in yellow corresponds to cross-β forming region. (B) Levels of ThT fluorescence were measured by spectroscopy as a function of time for protein samples corresponding to the native low complexity domain of TDP-43 (WT) or each of nine double proline substitution variants. Mean ± SEM; n > 4.

Obviously correlative effects of the double proline variants of the TDP-43 LCD were observed for their ability to overcome inhibition of GFP fluorescence by fragments 6 and 7 as assayed in bacterial cells (Figure 3), for their inhibitory effects upon the formation of liquid-like droplets as observed in test tube assays using purified proteins (Figure 4), and for their inhibitory effects upon assembly of the purified TDP-43 LCD into ThT-positive, cross-β polymers (Figure 5).

### Analysis of contiguously localized proline mutations within the TDP-43 LCD on the formation of nuclear speckles

The TDP-43 RNA binding protein resides primarily within the nuclear compartment of mammalian cells where it forms nuclear speckles (32). In order to determine whether self-association of the TDP-43 LCD might be required for its assembly into nuclear speckles, each double proline variant evaluated for phase separation (Figure 4), and formation of labile cross-β polymers (Figure 5), was introduced into lentivirus as a fusion protein linking GFP to the N-terminus of the full length TDP-43 protein. Viral infection allowed the GFP:TDP-43 fusion proteins to be conditionally expressed in HCT116 cells after doxycycline-mediated induction (Materials and Methods). Western blotting assays revealed that the nine variants were expressed at the same level as the fusion protein linking GFP to the native TDP-43 protein (Supplemental Data, Figure S3). The closely matched levels of expression of the various GFP:TDP-43 fusion proteins were approximately equivalent to that of endogenous TDP-43 within HCT116 cells.

Confocal microscopy was used to evaluate the morphological distribution of TDP-43 nuclear speckles. Cells expressing a GFP fusion protein linked to the native TDP-43 protein revealed roughly 30 GFP-positive speckles per nucleus (Figure 6). Cells expressing three of the double proline variants, designated F6-P3, F6-P4 and F6-P5, revealed a statistically significant reduction of roughly 75% in the number of GFP-positive nuclear speckles. Cells expressing the F6-P6 and F6-P7 variants revealed reductions of 40-50% in the number of GFP-positive nuclear speckles. Finally, cells expressing the F6-P1, F6-P2, F7-P6 and F7-P7 variants showed no statistically significant reduction in GFP-positive nuclear speckles relative to the number formed by the native TDP-43 protein.

**Figure 6.**
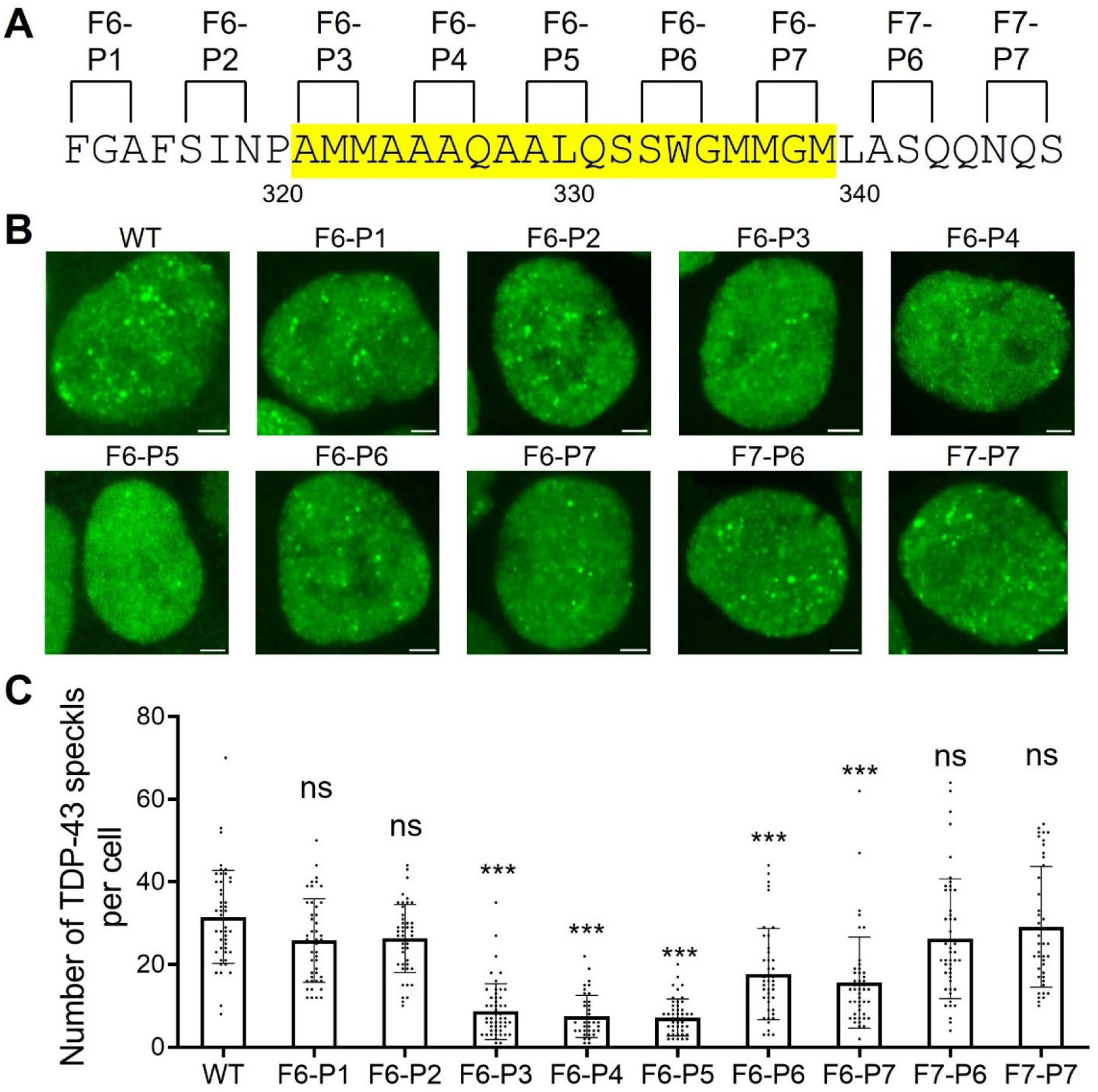
Effects of proline substitutions upon condensation of TDP-43 into nuclear speckles in cultured human cells. (A) GFP was fused onto the N-terminus of the full length TDP-43 protein containing either the native sequence of its low complexity domain (WT) or double proline variants thereof. Sequence highlighted in yellow corresponds to cross-β forming region. (B) Coding segments for each construct were moved from bacterial plasmids to lentivirus and used to infect HCT116 cells. Cell lines expressing each construct were selected for lentivirus-specified resistance to hygromycin (Materials and Methods), grown on cover slips and analyzed for nuclear GFP fluorescence by confocal microscopy. Scale bar: 2 μm. (C) 40 individual nuclei were analyzed for each cell line to quantify the number of GFP-positive nuclear speckles per cell. Mean ± SD; ***p < 0.001; ns, no significance; one-way ANOVA.

In summary, correlatively similar patterns of effect of double proline mutation were observed for: (i) relief of fragment 6- and fragment 7-mediated suppression of GFP fluorescence (Figure 3); (ii) formation of phase separated liquid-like droplets (Figure 4); (iii) formation of labile cross-β polymers (Figure 5); and (iv) assembly of TDP-43 into GFP-positive nuclear speckles in cultured HCT116 cells (Figure 6). In combination, these data confirm that the TDP-43 LCD achieves self-association via the formation of a labile cross-β structure localized to an evolutionarily conserved region of roughly twenty amino acid residues.

### Two related aliphatic alcohols differentially melt nuclear speckles formed by TDP-43

The GFP-suppressing segment of the TDP-43 LCD co-localizes with a region where differences in protein binding by two related aliphatic alcohols have been mapped. Solution NMR studies have shown that 1,6-hexanediol (1,6-HD) causes significantly more pronounced chemical shifts to backbone carbonyl oxygen atoms than 2,5-hexanediol (2,5-HD) in the region of the TDP-43 LCD bracketed by alanine residue 321 and leucine residue 340 (33).

Differences in the capacity for sequestration of backbone hydrogen bond acceptors explain the enhanced potency of 1,6-HD, relative to 2,5-HD, for melting phase separated liquid-like droplets and labile cross-β polymers formed by the TDP-43 LCD (33). This form of protein:protein interaction requires the zippering of a contiguous set of interstrand hydrogen bonds between peptide amino groups and carbonyl oxygens. When the terminal alcohol groups of 1,6-HD become hydrogen bonded to backbone carbonyl oxygen atoms within this evolutionarily conserved region of the protein, self-association is chemically inhibited.

Knowing that aliphatic alcohols bind to carbonyl oxygen atoms of the polypeptide backbone of LCDs (33, 34), our mechanistic interpretation of TDP-43 self-association predicts that these pharmacological agents should melt nuclear speckles formed by TDP-43. This expectation is based upon the assumption that the partitioning of TDP-43 into intranuclear condensates is dependent upon the ability of its LCD to self-associate.

Indeed, this concept further predicts that nuclear speckles containing the TDP-43 protein should be dissolved more readily by 1,6-HD than its 2,5-HD regioisomer. Differences in the dissociative activities of the two aliphatic alcohols have been observed in living cells for a variety of nuclear and cytoplasmic puncta not surrounded by investing membranes, as well as for the integrity of five different types of intermediate filaments (27). That 1,6-HD is more effective in melting self-associated LCDs than 2,5-HD has been attributed to geometrical differences in the positions of the two alcohol groups along the hexameric carbon chain of each chemical. The spacing of OH groups in 1,6-HD better matches the distance separating contiguous carbonyl oxygens along the polypeptide backbone than 2,5-HD. As such, 1,6-HD is better suited, relative to 2,5-HD, for forming two concomitant hydrogen bonds to adjacent carbonyl oxygen atoms along the polypeptide backbone (33, 35).

To test the effects of aliphatic alcohols on nuclear speckles formed by TDP-43, cultured HCT116 cells expressing the full-length TDP-43 protein fused at its N-terminus to GFP were exposed to 8% levels of either 1,6-HD or 2,5-HD for a period of five minutes (Materials and Methods). As shown in Figure 7, the number of nuclear speckles containing the GFP:TDP-43 fusion protein was reduced by roughly 75% following five minutes of exposure to an 8% concentration of 1,6-HD. Exposure of the same cells to an equivalent concentration of 2,5-HD reduced speckle number by only 30% relative to vehicle-treated cells. These same differences in the melting of TDP-43 nuclear speckles by the same two aliphatic alcohols have been reported by Fawzi, Shorter and colleagues (32). As such, two independent studies offer concordant observations pertinent to the chemical underpinnings of TDP-43 self-association within the nuclei of cultured mammalian cells.

**Figure 7.**
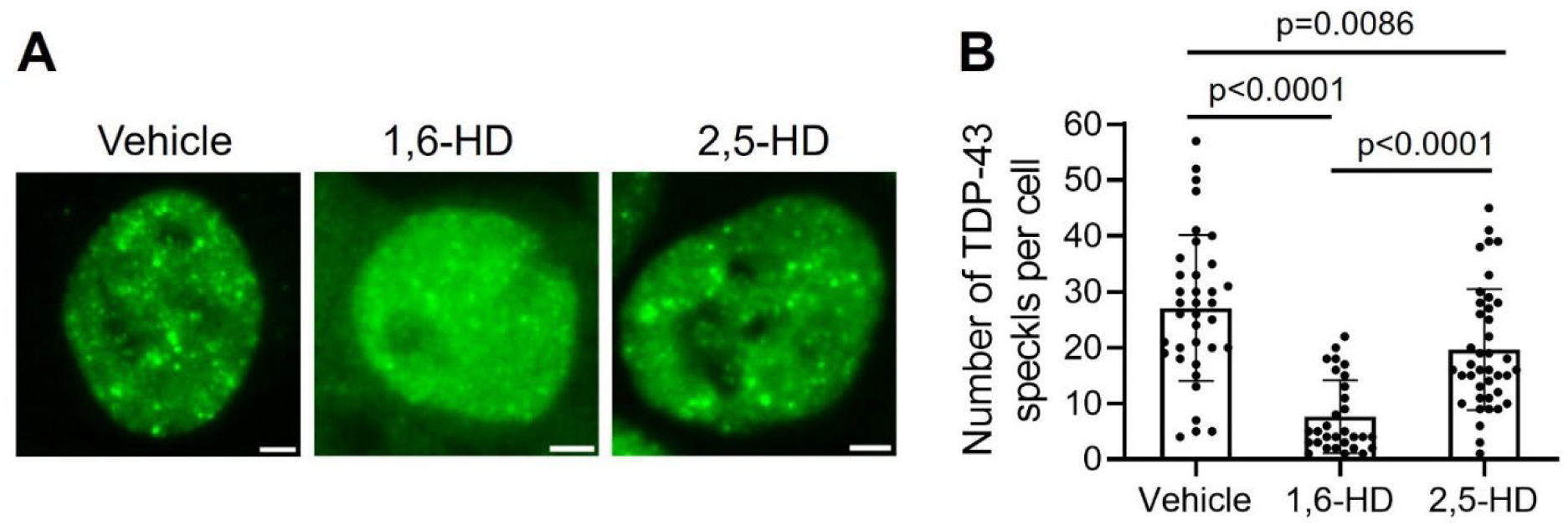
Effects of two regioisomeric alcohols on nuclear speckles composed of TDP-43. (A) Confocal microscopic images of HCT116 cells expressing a lentivirus-encoded GFP:TDP-43 fusion protein. Cells were grown on cover slips and exposed for 5 minutes to vehicle alone (left), an 8% solution of 1,6-hexanediol (middle), or an 8% solution of 2,5-hexanediol (right). Scale bar: 2 μm. (B) Focal plane images of 30 individual nuclei were analyzed for each culture condition to quantify the number of GFP-positive nuclear speckles. Mean ± SD; one-way ANOVA.

## Discussion

Here we describe a simple method for mapping the region of a protein domain of low sequence complexity required for its self-association, phase separation and partitioning into nuclear puncta. The method we describe should facilitate analysis of any LCD that relies upon homotypic self-association to perform its biological function.

We begin our discussion by asking why several short fragments of the TDP-43 LCD interfere with GFP-fluorescence? The slow-to-fold GFP protein is structurally organized as a barrel-like ensemble of eleven β sheets (36, 37). The final β strand required to complete folding of the protein is located at the very C-terminus of GFP. Insertion of this final β strand represents a slow, co-translational event that is rate-limiting to GFP folding (38). We speculate that C-terminal juxtaposition of an additional, β strand-prone segment somehow impedes the completion of GFP folding. Owing to the high level of bacterial expression of the GFP:TDP-43 fusion proteins studied herein, we recognize that observed impediments to GFP fluorescence may involve self-association of certain test protein fragments. This interpretation may explain why proline substitutions that interfere with the formation of cross-β structural assemblies, as shown in Figures 4 and 5, also overcome the suppression of GFP fluorescence as shown in Figure 3.

Turning from methodology to investigational focus, our findings are interpretable according to a simple concept of TDP-43 LCD self-association. As reported previously, the TDP-43 LCD self-associates via the formation of a labile, cross-β structure localized to a small region of high evolutionary conservation (16, 24). The GFP suppression screens described in Figures 1 and 2 map inhibition to the same region of the TDP-43 LCD known to mediate self-association and phase separation. Moreover, the observed GFP inhibitory activity was overcome by proline mutations localized to the cross-β forming region of the LCD. Concordance was observed between proline variants that overcame GFP inhibition (Figure 3) with those that inhibited condensation of liquid-like droplets (Figure 4), inhibited assembly of cross-β polymers (Figure 5), and attenuated formation of TDP-43 positive nuclear speckles (Figure 6).

This structure-based interpretation of LCD self-association contrasts with a distinctly different concept having emerged from computer-assisted simulations of LCD phase separation (39, 40). The latter, “stickers-and-spacers” concept is incongruous to the labile cross-β model for LCD self-association, phase separation and biological function in three ways: (i) it posits that LCD self-association is achieved in the absence of molecular structure; (ii) it assigns no role for the polypeptide backbone in the annealing of polypeptide strands; and (iii) it assumes that the interactions required for self-association cannot be sub-localized by either experimentation or computation. We are currently unable to reconcile the data reported herein with computer-simulated interpretations of LCD self-association.

Why is it useful to know where the small, cross-β forming regions mediating self-association and phase separation map within larger LCDs? First, this understanding helps explain how single residue, disease-causing mutations interfere with LCD function. As an example, by localizing the cross-β forming region that allows the head domain of the neurofilament light (NFL) protein to weakly self-associate in a manner abetting filament assembly, we can understand why virtually any mutational variation of either proline residue 8 or proline residue 22 causes Charcot Marie-Tooth disease (15). These proline residues insulate either side of the localized, cross-β forming region of the NFL head domain. Mutational change of either of these proline residues to any of the other 19 amino acids introduces an additional peptide NH group to the polypeptide backbone. As such, alteration of proline residues proximal to cross-β forming regions can be understood to enhance the avidity of self-association by addition of an interstrand hydrogen bond (16). As a consequence, mutational change of proline residues can cause evolutionarily balanced forms of protein:protein interaction to be forced out of tune.

A second example of the value of knowing where cross-β interactions map within LCDs also derives from studies of the disordered head domains of intermediate filament (IF) proteins. IF head domains are phosphorylated during mitosis, thus triggering filament disassembly and allowing cultured cells to round up during mitosis (41, 42). The sites of protein kinase A-mediated phosphorylation of the desmin IF protein map to the cross-β forming region of its head domain. Moreover, phosphorylation of the desmin head domain reverses self-association and phase separation by weakening labile, cross-β interactions (15). As such, the biological phenomenon of IF disassembly and consequent change in cell shape during mitosis is now understood at a mechanistic level.

A third justification favoring the reductionist mapping of cross-β forming regions within LCDs relates to the aspired evolution of this science from test tubes to computers. Development of AlphaFold capabilities of protein structure prediction required a foundation consisting of thousands of structures deduced by X-ray crystallography, NMR spectroscopy and cryo-electron microscopy (43). The methods described herein will simplify the mapping of cross-β forming regions within larger LCDs, thereby accelerating the conversion of this science from lab bench to laptop.

We close with a thread of logic connecting chemistry to biology. Thousands of studies published over the past decade have reported evidence confirming that LCD self-association is of widespread importance to the dynamics of cell morphology (21, 44, 45). A cursory evaluation of the Google Scholar database tagged by the key words of *phase separation*, *1,6-hexanediol* and *low complexity domain* reveals more than 400 of these studies as having demonstrated the melting of nuclear or cytoplasmic structures in response to 1,6-hexanediol (Supplemental Data, Figure S4). We propose that if an intracellular structure is dissolved more readily by 1,6-hexanediol than 2,5-hexandiol, as demonstrated herein for nuclear speckles composed of TDP-43 (Figure 7), the structure of interest will be reliant upon labile cross-β interactions. This logic, coupled with the experimental method of LCD dissection described herein, offers a conceptual pathway for demystifying many forms of dynamic cell organization.

## Materials and Methods

### Plasmid construction

DNA sequences encoding the TDP-43 low complexity domain (LCD, residue 262-414) were sub-cloned into a pHis-parallel vector with an N-terminal His tag. DNA fragments encoding 30 amino acid residues (fragments 1 through 12) from the TDP-43 LCD were sub-cloned into pHis-parallel-GFP-GDEVD with BamHI and XhoI restriction endonuclease sites. Proline variants of His-TDP-43 LCD, GFP-TDP-43 fragment 6 (F6) and GFP-TDP-43 fragment 7 (F7) were generated by site-directed mutagenesis.

### Protein expression and purification

His-TDP-43 LCD WT and mutated variants were expressed in *E. coli* BL21(DE3) cells at 37 °C, induced with 1 mM IPTG for 3 hours. The harvested cell pellets were lysed in a buffer containing 50 mM Tris (pH 7.5), 6 M guanidine-HCl, and 20 mM imidazole. Lysis was performed via sonication (3 minutes total, alternating 10 seconds on and 30 seconds off). Cell debris was removed by centrifugation at 25,000 RPM for 30 minutes. The clarified supernatant was loaded onto a gravity-fed column packed with Ni-NTA resin (Gold Bio). The resin was washed with the lysis buffer to remove non-specifically bound proteins. Target proteins were eluted using a buffer containing 50 mM Tris (pH 7.5), 6 M guanidine-HCl, and 300 mM imidazole. Eluted proteins were concentrated to 1 mM and subjected to buffer exchange prior to use. Buffer exchange was performed by dialysis against a storage buffer containing 10 mM Tris (pH 7.5), 6 M guanidine-HCl, and 1 mM EDTA using Slide-A-Lyzer MINI dialysis units (Thermo Scientific, 69552) at 4 °C. Protein concentrations were determined using both a Nanodrop spectrophotometer and a BCA assay. For droplet formation assays and Thioflavin T (ThT) fluorescence kinetic assays, protein concentrations were adjusted to 600 μM using storage buffer.

### Droplet formation and OD_600nm_ measurements

His-TDP-43 LCD WT and its variants (600 μM in 6 M guanidine-HCl) were diluted 30-fold into droplet formation buffer as indicated using a multichannel pipette. A 40 μL aliquot of the resulting solution was transferred into a 384-well plate (Greiner Bio-One, 781091) using the same pipette. After a 5-minute incubation, absorbance at 600 nm was measured using a microplate reader. For droplet formation imaging, samples were incubated for 1 hour, and images were acquired using the Bio-Rad ZOE Fluorescent Cell Imager.

### Thioflavin T (ThT) fluorescence kinetic assays

His-TDP-43 LCD WT and its variants (600 μM in 6 M guanidine-HCl) were diluted into ThT buffer containing 50 mM MES (pH 6.8), 150 mM NaCl, 1 mM EDTA, 40 μM ThT, and 0.05% NaN₃. The final protein concentration was adjusted to 20 μM. All ThT samples were dispensed into a 384-well microplate (Greiner Bio-One, 781091) with a volume of 40 μL per well. The plate was sealed with a foil film to prevent evaporation. ThT fluorescence was monitored at room temperature with an excitation wavelength of 450 nm and an emission wavelength of 485 nm. The master plate was shaken for 10 seconds before each reading of fluorescence to ensure sample uniformity.

### GFP suppression assays

Plasmids encoding N-terminal GFP-tagged TDP-43 fragments and mutated variants thereof were transformed into BL21(DE3) competent cells via heat shock following standard protocols. Transformed cells were incubated at 37 °C in a shaker for 40 minutes. Cells were subsequently inoculated into 96 DeepWell plates (Fisher, 12566612) containing 0.5 mL of LB culture medium supplemented with 100 μg/mL ampicillin and 40 μM IPTG. Plates were sealed with Constar 6570 sealing tape and incubated overnight at 37 °C with shaking. Following incubation, the cells were harvested by centrifugation, washed once with PBS, and resuspended in 1 mL of PBS. GFP fluorescence intensity expressed in living *E. coli* cells was measured alongside turbidity, as both values exhibit a linear correlation under appropriate cell densities. *E. coli* suspensions were serially diluted followed by measurements of both turbidity (absorbance at 600 nm) and GFP fluorescence intensity (excitation/emission = 488 nm/513 nm) using transparent 96-well plates (NEST, 701011, 100 μL per well) and a microplate reader. The resulting data were fitted to linear regression curves using GraphPad. The slopes of the WT and double proline variant curves were recorded and normalized to the WT value for comparison.

### Cell culture

HCT116 cells were a gift from Dr. Deepak Nijhawan. HCT116 cells were cultured in the Dulbecco’s modified Eagle’s medium (DMEM) supplemented with 10% fetal bovine serum, 2 mM L-glutamine. All cells were incubated at 37 °C in a humidified atmosphere of 95% air and 5% CO_2_.

### Lentivirus production and generation of inducible cell lines

Full-length GFP-TDP-43 WT and proline mutants thereof were cloned into a doxycycline-inducible pSPCTRE-Hygro vector using the AscI and BamHI restriction sites. LentiX293T cells were co-transfected with psPAX2, pMD2.G, and pSPCTRE-Hygro-GFP-TDP-43 plasmids. After 24 hours, the culture medium was replaced. At 72 hours post-transfection, the medium was collected and filtered through a 0.45 μm syringe filter (Millipore, SLHV033RB). The resulting lentiviruses were used to infect HCT116 cells, which were subsequently selected using 200 μg/mL hygromycin.

### Protein extraction and western blotting

Inducible cell lines expressing GFP-TDP-43 full-length WT and double proline variants thereof were cultured in 6-well plates. After 24 hours of incubation, doxycycline was added to the medium at a final concentration of 3 ng/mL. Following 48 hours of induction, the cells were washed twice with PBS and lysed in RIPA buffer (30 mM Tris, pH 7.4, 150 mM NaCl, 1% IGEPAL CA-630, 1% sodium deoxycholate, 0.1% SDS) supplemented with a protease inhibitor cocktail (Roche) and Benzonase Nuclease (Novagen). Lysis was performed on ice for 30 minutes. Cell debris was removed by centrifugation at 21,000 × g for 30 minutes at 4 °C. Protein concentration of the lysates was determined using a BCA Protein Assay Kit (Thermo Fisher Scientific). For each cell line, 10 μg of total protein was resolved by SDS-PAGE. Proteins were transferred to a membrane and immunoblotted with primary antibodies, including rabbit anti-N-terminal TDP-43 (Proteintech, 10782-2-AP) and mouse anti-GAPDH (EMD Millipore, C87727). Secondary antibodies used were HRP-conjugated goat anti-rabbit and HRP-conjugated goat anti-mouse (Bio-Rad). Chemiluminescence detection was performed using the ECL Western Blotting Substrate (Bio-Rad), and images were captured with a ChemiDoc Touch Imaging System (Bio-Rad).

### Immunocytochemistry and confocal imaging

Inducible cell lines expressing GFP-TDP-43 full-length WT and double proline mutants thereof were cultured in 4-well chambers (BD Falcon, 354104) and incubated with 3 ng/mL doxycycline for 48 hours. After removing the culture medium, the cells were washed twice with warm PBS and fixed with 4% PFA for 15 minutes. Following fixation, cells were washed twice with PBS and mounted using Vectashield mounting medium supplemented with DAPI dye (Vector, H1200). Confocal images were acquired using a Zeiss LSM880 microscope equipped with a 63× oil immersion objective, capturing 0.5 μm Z-stacks at 3.5× zoom. Image dimensions were set to 1024 × 1024 pixels. For hexanediol-treated cells, GFP-TDP-43 WT-expressing cells were cultured in poly-D-lysine-coated chambers (Gibco, A3890401), incubated with 3 ng/mL doxycycline for 48 hours, and subsequently treated with 8% 1,6-hexanediol (Sigma-Aldrich, 240117) or 8% 2,5-hexanediol (Sigma-Aldrich, H11904) for 5 minutes before washing and fixation. Confocal images were acquired using parameters described above.

Analysis of TDP-43 nuclear speckles was conducted using Fiji software. Confocal images of GFP-positive nuclei were processed with maximum intensity Z-projection and smoothed. Single GFP-positive nuclei were extracted, and their average fluorescence intensity was calculated by using Image → Adjust → Threshold → Method: Li → Auto → Analyze → Analyze Particles modules. Speckles were identified as structures with fluorescence intensity exceeding 1.6 times the average intensity. Adjacent speckles were separated using Process → Binary → Watershed, and speckle count was automatically determined with Analyze → Analyze Particles.

### Quantification and statistical analysis

Statistical parameters including the definitions and values of n, distributions and deviations are reported in Figures and Figure Legends. Measurements of statistical significance in this were determined by One-way ANOVA. Statistical analysis was performed in GraphPad.

## Data, Materials, and Software Availability

All data presented in this study are available within the Figures and in SI Appendix. All study data are included in the article and/or SI Appendix.

## ACKNOWLEDGMENTS

This work was supported by an anonymous donor (funds to S.L.M.), the National Institute of General Medical Science (grant GM130358 to S.L.M.)

## Supplementary Data Figures and Legends

**Supplementary Data Figure S1.**
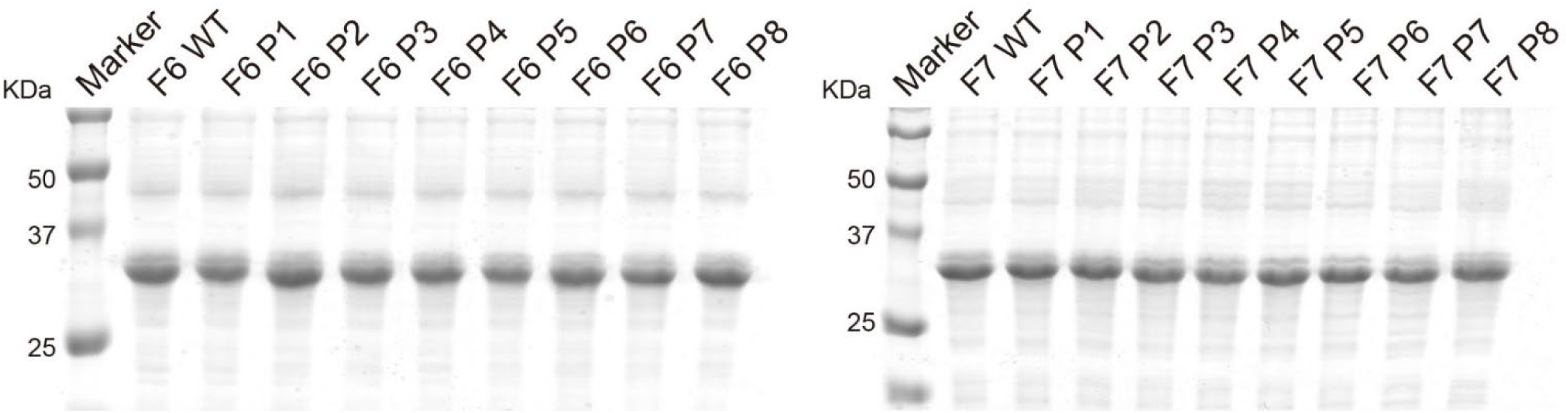
Coomassie stained SDS-PAGE gels used to visualize fusion proteins linking GFP to either fragment 6 (left panel) or fragment 7 (right panel) of the TDP-43 low complexity domain. In addition to GFP fusions linked to the native sequences of fragment 6 (F6 WT) and fragment 7 (F7 WT) of the TDP-43 low complexity domain, gel images show GFP fusions linked to double proline variants of each fragment as described in Figure 3 of main text. Molecular weight markers are shown in the left lane of each SDS gel.

**Supplemental Data Figure S2.**
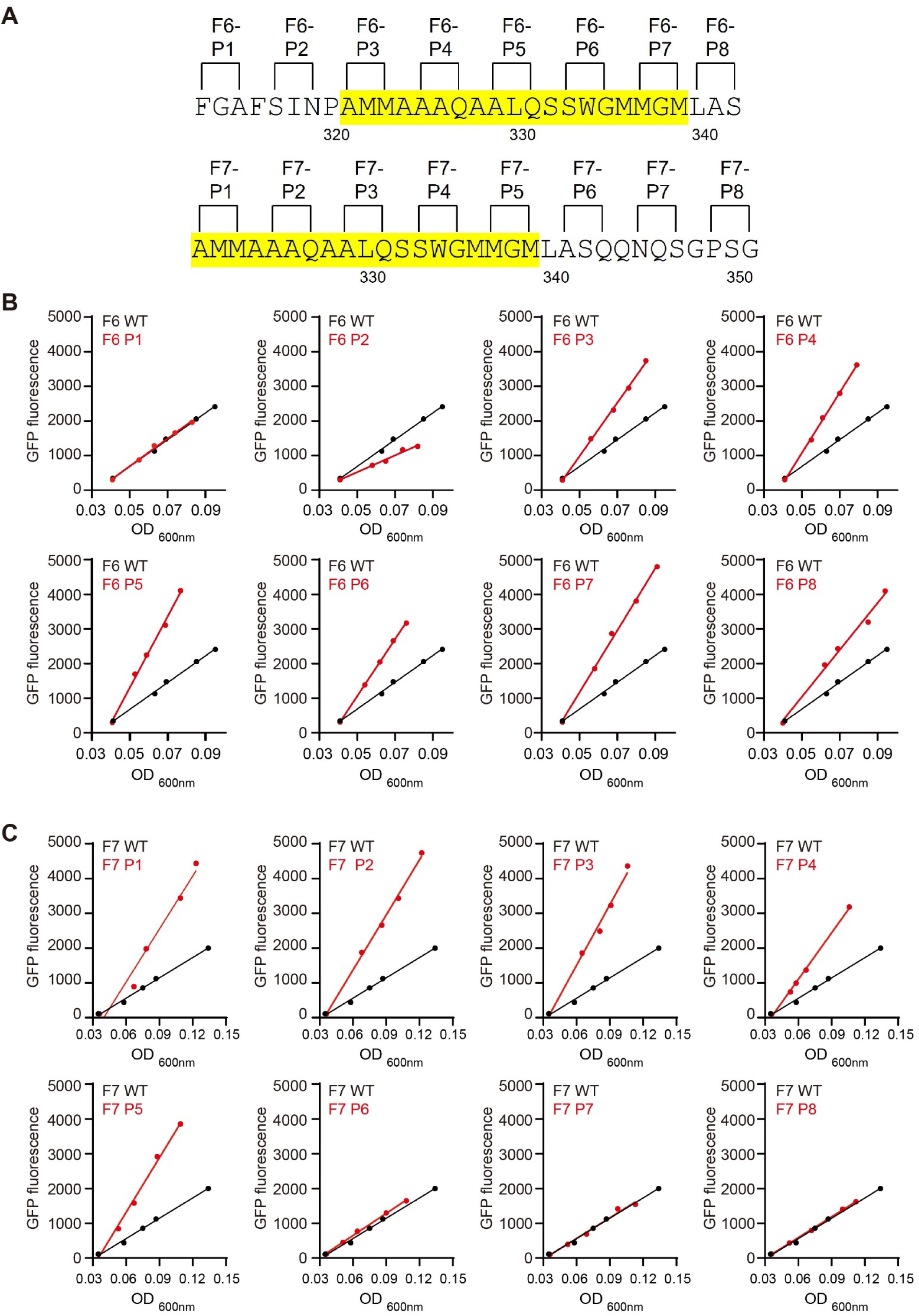
Measurements of GFP fluorescence from fusion proteins of the TDP-43 30 residue fragments and double proline variants thereof. **(A)** Sequences of fragment 6 (above) and fragment 7 (below) of the TDP-43 low complexity domain with schematic designation of locations of double proline variants. Sequences highlighted in yellow correspond to cross-β forming region. **(B and C)** Measurements of increases in OD_600nm_ as a function of cell density (X axis) plotted against measurements of increases in GFP fluorescence as a function of cell density (Y axis). Black lines for each graph of panel **(B)** describe measurements for cells expressing GFP fused to fragment 6 of the TDP-43 low complexity domain. Black lines for each graph of panel **(C)** describe measurements for cells expressing GFP fused to fragment 7 of the TDP-43 low complexity domain. Red lines of both panels **(B and C)** describe measurements for individual double proline variants. Slope calculations for each double proline variant were used for data shown in Figure 3.

**Supplementary Data Figure S3.**
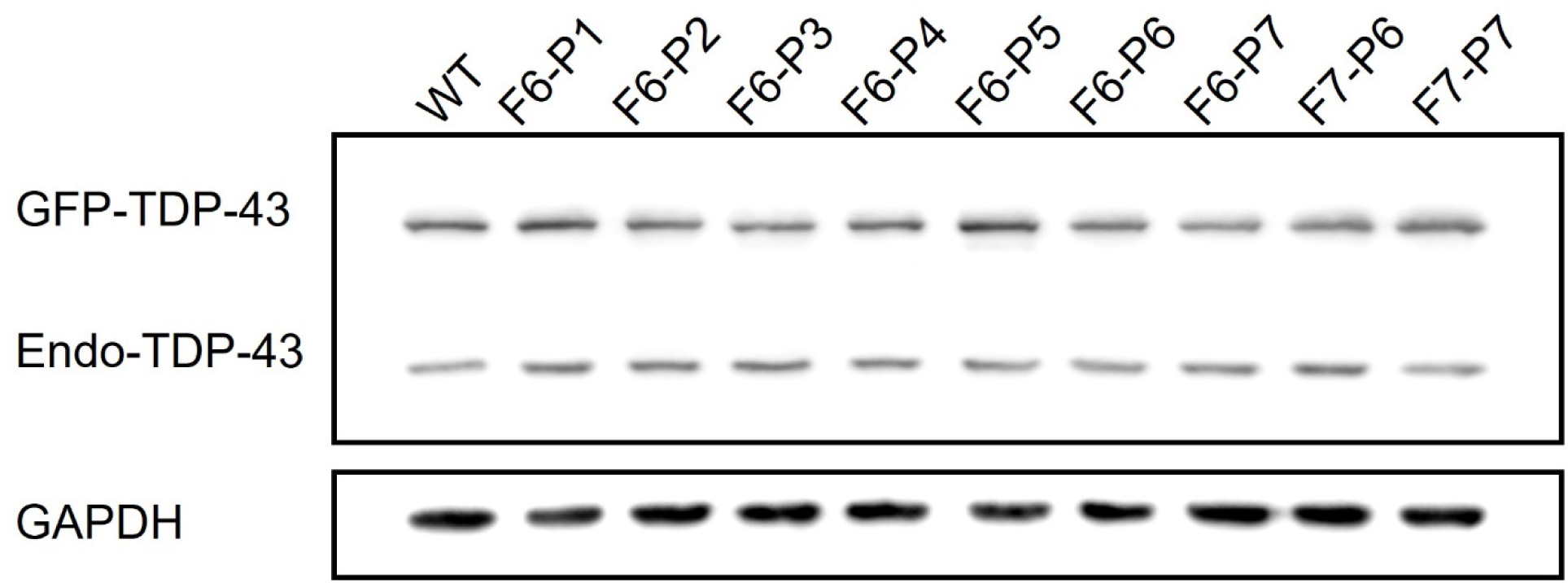
Western blot assays for the expression of lentivirus-encoded GFP:TDP-43 fusion protein in HCT116 cells. Top panel shows Western blot of cell lines expressing GFP fusion proteins linked to either the full length, native TDP-43 protein, or to the full length TDP-43 protein bearing designated double proline variants. The antibody used for top Western blot was specific to the native TDP-43 protein and thus visualized both the exogenous, virus-encoded GFP:TDP-43 fusion protein (GFP-TDP-43) or the endogenous TDP-43 protein of HCT116 cells (Endo-TDP-43). Bottom panel shows Western blot of same samples as assayed in top panel, yet probed with an antibody specific to the GAPDH enzyme endogenous to HCT116 cells.

**Supplementary Data Figure S4.**
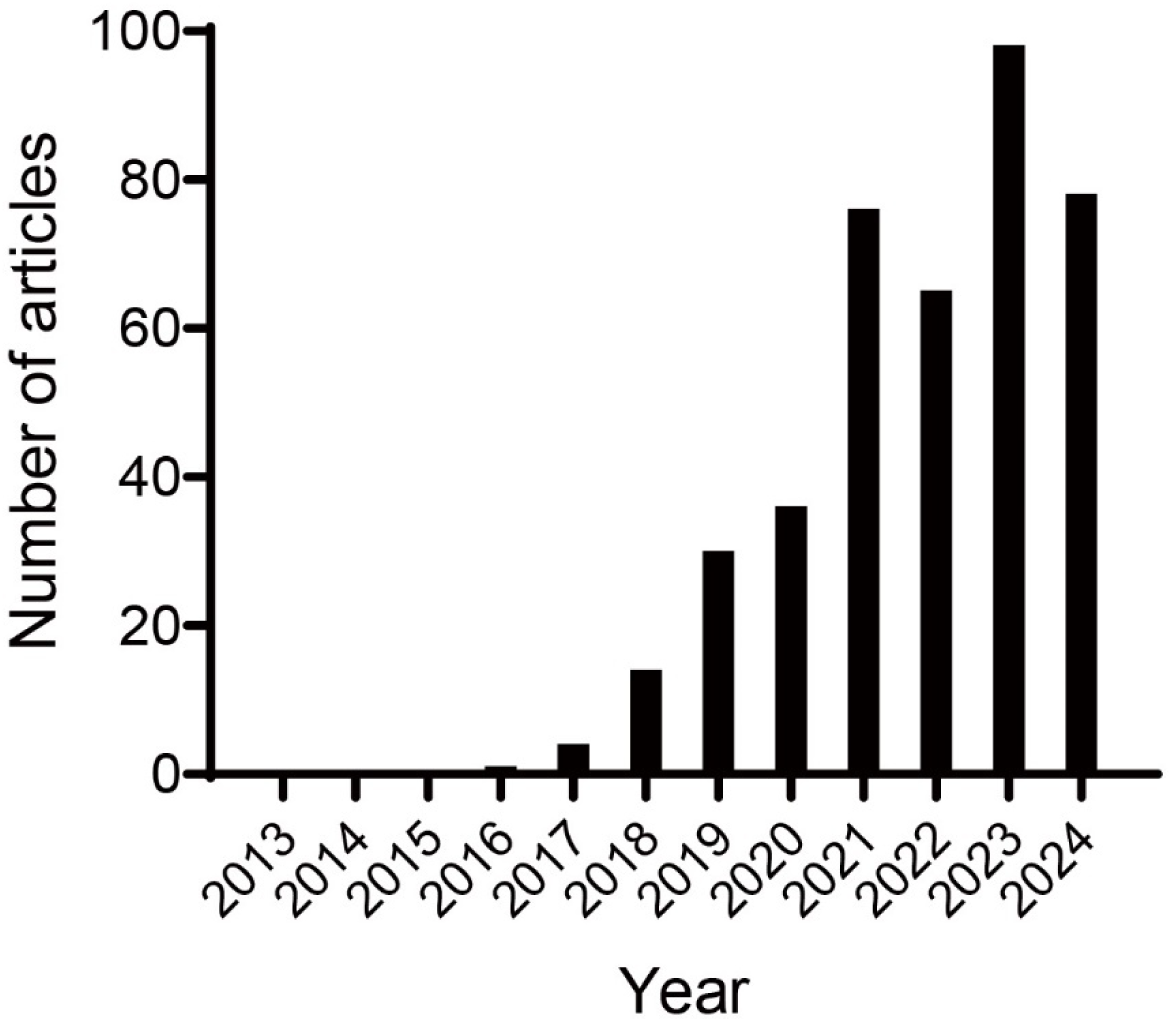
Google Scholar reported research articles between 2013 and 2024 as identified by all of three key words, *phase separation, 1,6-hexanediol* and *low complexity domain*.

